# Transient signaling of free ADP-ribose monitored with an intracellular biosensor

**DOI:** 10.64898/2026.02.06.704453

**Authors:** Shivansh Goyal, Vanna Nguyen, Scott N Lyons, Tyler L Dangerfield, Wanjie Yang, Ciara Fields, Hsin-Ru Chan, Yazhini Mahendravarman, Lorren Cantu, Nesha Rubin, Kelton Deloney, Kenneth A Johnson, Xiaolu A. Cambronne

**Affiliations:** Department of Molecular Biosciences, University of Texas at Austin, Austin TX 78712 USA

## Abstract

We have developed a biosensor enabling the dynamic, compartmentalized, and longitudinal measurements of intracellular ADP-ribose (ADPR) in live cells. Free ADPR is a critical signaling metabolite derived from nicotinamide adenine dinucleotide (NAD^+^). As an agonist for Transient Receptor Potential Melastatin 2 (TRPM2), ADPR levels can regulate immune responses during infection, as well as nociception and adjustment of core body temperature. The study of ADPR signaling has been limited, however, by a lack of methods to measure this metabolite in situ. Using the biosensor and its paired non-responsive control, we determine that intracellular ADPR accumulation was transient and tunable. We found that basal concentrations were in the nanomolar range and could be stimulated ∼30-fold to activate TRPM2. We identified that TRPM2 activation, measured by calcium influx, required an intracellular ADPR threshold concentration between 2 – 4 µM at physiological temperature. We observed that the timing of the ADPR rise coincided with TRPM2 activation, thus providing support for ADPR fluctuations being a critically regulated aspect for channel activation. Notably, transient fluctuations of ADPR were not accurately reflected by measurements of intracellular NAD^+^ loss or calcium levels.

**Significance Statement:** We have developed a unique real-time biosensor for free ADP-ribose that is tuned to physiological concentrations and capable of intracellular measurements in individual cells. Using a calibrated system we determined that concentrations and timing of induced intracellular ADPR aligned with the thermosensitive TRPM2 activity. The data support ADPR as a critical component whose intracellular levels are regulated to control TRPM2 channel opening in cells.

## Introduction

Free ADP-ribose (ADPR) is a metabolic second messenger derived from the cleavage of nicotinamide adenine dinucleotide (NAD^+^). By association with the enzymes that produce and use the signaling molecule, free ADPR signaling has been linked to immunomodulation, fever and oxidative stress responses, cancer, diabetes, neurodegeneration, and neuropathy (1–15).

Elucidating how cells use and regulate levels of intracellular free ADPR has been challenging due to a lack of direct measurements in intact cells. Enzymes that catalyze ADPR production and turnover are spatially organized and thought to locally regulate the abundance of this signaling molecule (16). ADPR can artefactually arise during the extraction process and is a common contaminant in commercially available cADPR and adenine dinucleotides (14, 17–19). In short, there are unknowns about the regulation of free ADPR that challenge our understanding of its targets and mechanisms of action (16, 19, 20).

Transient Receptor Potential Melastatin 2 (TRPM2) is a ligand-gated cation channel that predominantly functions at the plasma membrane (21–25). Convergent structural, biochemical, and electrophysiological data have established that free ADPR (and not NAD^+^, NAADP, or cyclic-ADPR) (17–19) is an endogenous agonist for calcium-bound TRPM2 under physiological temperatures (26). While remaining strictly ligand-gated, TRPM2 integrates extracellular cues of calcium availability and temperature with levels of intracellular ADPR ligand for full channel opening. Additional reported roles for ADPR include activating intracellular ryanodine receptors (27) and extracellular P2Y1 receptors (5, 28, 29) and for substrate-level ATP production in the nucleus (30, 31). ADPR has been further reported to bind to Complex I in mitochondria (32–35) and to cytosolic glycolysis enzymes (36).

Consequently, ADPR is a critical contributor to cellular processes and study of its roles requires methods to quantitatively monitor ADPR dynamics in situ. Previously, a Förster resonance energy transfer (FRET)-based assay was developed that used cysteine labelling on a catalytically inactivated NUDT9 protein via maleimide click chemistry (37). However, its applicability has been largely limited for assays in vitro. Molecular probes such as *Af*1521 and engineered *Af*1521 Fc fusion proteins can bind ADPR but do not distinguish the free from covalently bound molecules in situ (38, 39). Similarly, existing genetically encoded sensors that detect poly-ADPR chains are used for monitoring PARP activity but do not detect free ADPR molecules (40–42).

We have developed a genetically encoded single fluorescent protein biosensor that selectively monitors free ADPR with ratiometric intensity changes. We determined that ADPR signaling is transient and cannot be accurately reflected by proxy measurements of NAD^+^ or free calcium fluctuations. We found that basal ADPR levels were held < 1 μM and only accumulations > 2 μM elicited TRPM2 activation under physiological conditions, thus establishing thresholds of intracellular ADPR levels regulating TRPM2 channel opening.

## Results

### A selective and dynamic biosensor for free ADPR

We recognized that bacterial transcriptional repressor NrtR de-repressed its targets in response to direct binding of intracellular ADPR (43). Structural analyses of *Shewanella oneidensis* NrtR showed multiple ADPR-dependent allosteric changes enabling its release from DNA, including a Nudix switch element that transformed from an alpha helix into an extended state (Fig. S1A) (44). Insertion of circularly permutated Venus fluorescent protein (cpVenus) within the nudix switch at Q118 created a candidate biosensor, and we introduced additional point mutations D122N, K435A and S436D to improve pH resistance, brightness, and intracellular solubility (Fig. S1B).

We determined that the ADPR biosensor exhibited a ∼3-fold decrease in fluorescence intensity in vitro in the presence of saturating ADPR concentrations (Fig. 1A). We created a non-responsive fluorescent variant (R98E) to serve as a negative control, and this non-binding variant showed no significant fluorescent changes between tested 100 pM - 1 mM free ADPR (Fig. 1A). We found that the biosensor was consistent for reporting relative ADPR fluctuations between pH 6.5 to 8.0 (Fig. 1B) but that its basal brightness was 1.65-fold dimmer at pH 6.5 compared to pH 8.0 (Fig S1C). This pH-dependent change in brightness, nevertheless, was improved from the original 5.5-fold change in brightness of its fluorescent protein cpVenus across this range (Fig S1C). Increasing temperature (tested up to 42 °C) was another determinant that dimmed the basal brightness of the biosensor (Fig. 1C). The non-responsive R98E control was similarly affected by temperature, and when incorporated for normalization, resulted in stabilized ADPR measurements between 25°C – 42°C (Fig. S1D). In vitro stopped-flow kinetic measurements revealed dynamics consistent with a single site binding (Fig. 1D). The data were fit using nonlinear regression based on numerical integration of the rate equations, and we observed high confidence in the global data fit (Fig. 1E) with the proposed on-step binding mechanism 𝐸 + 𝑆 ⇌ 𝐸𝑆 and k_on_ =1.42 µM ± 0.02 µM, k_off_ = 0.84 µM ± 0.014 µM (Fig. 1F). An independent plotting of experimentally replicated datapoints revealed congruent values (Fig. S1E, S1F). Analysis of the experimentally solved binding pocket in NrtR revealed that ADPR bound in a linear conformation (Fig. 1G), suggesting selectivity for ADPR over cyclic-ADPR. In agreement, the sensor responded to ADPR and physiological analogs (2’-deoxy-ADPR, 2’/3’-O-acetyl-ADPR, and phospho-ADPR), but it did not respond to cyclized ADPR nor other tested nucleotides at their expected physiological concentrations (Fig. 1H); the R98E control did not respond to any tested analyte (Fig. S1G). Notably, ADPR, acetyl-ADPR, phospho-ADPR and 2’deoxy-ADPR are endogenous agonists of TRPM2 (45, 46), and both ADPR and phospho-ADPR are endogenous ligands of NrtR, reflecting the substrate accommodation of many Nudix-like domains found in natural enzymes (18, 44, 47).

**Figure 1.**
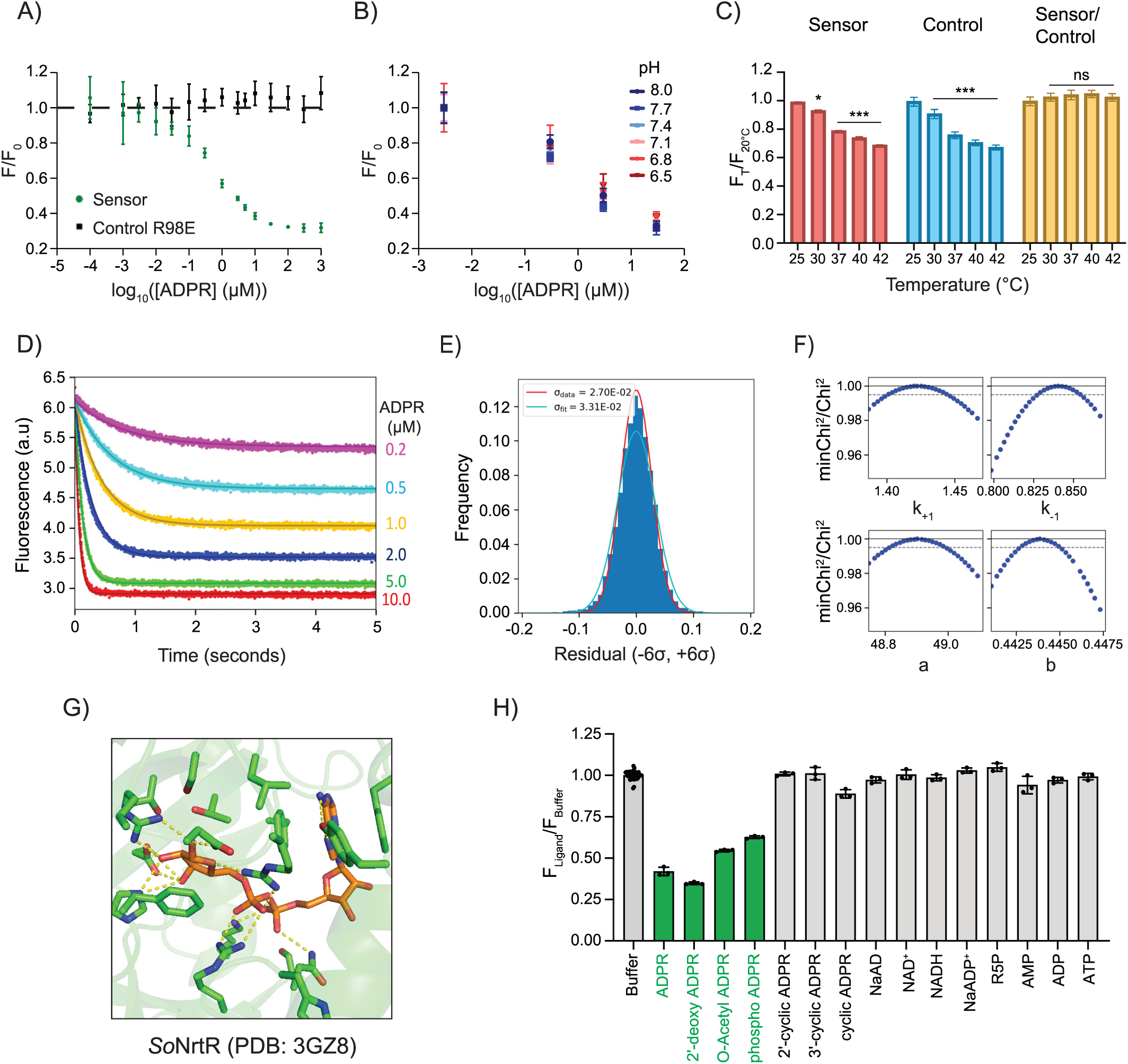
Characterization of a biosensor for free ADPR. **A)** In vitro dose responses of recombinant Sensor and R98E control at 20°C and pH 7.4, normalized to 0 μM ADPR (F_0_). **B)** ADPR-dependent sensor responses at indicated pH, 20°C, normalized to lowest ADPR concentration (0.003 μM, F_0_). **C)** Temperature effect on fluorescence of sensor and R98E control. **D)** Global fitting of stopped flow ADPR binding data. ADPR concentrations tested are listed to the right of the panel. **E)** Histogram of residuals (dark blue) calculated from the simulated global fit (red line, σ ∼0.02) compared to an ideal fit (cyan, σ ∼0.03). **F)** Confidence contour plots showing upper and lower limits based on min χ^2^ threshold of 0.99 (dotted line), corresponding to the 95% confidence interval. **G)** An ADPR molecule binds to *So*NrtR in extended conformation (PDB ID: 3GZ8). **H)** Effects of different ligands on the ADPR sensor. Ligands were tested at 10 µM, except for ADP, AMP, ATP at 500 μM, *n =* 39 without and *n =* 3 with ligands. Fluorescence data in A-C and H were measured using a fluorimeter at emission 520 ± 2 nm following excitation at 508 ± 2 nm, and plotted as mean ± SD, *n* = 3.

### Quantitative subcellular measurements of ADPR

To improve the precision of in-cell measurements, we fused the sensor with far-red fluorescent protein mCardinal (48) to normalize for variations in sensor expression (Fig. 2A). We found that ratiometric Green/Red emissions resulted in more consistent in-cell measurements compared to Green-only emissions. We quantified this as a 12-fold reduction in the heterogeneity in fluorescence (Interquartile range/Geometric Mean) across a population of cells expressing the sensor at low-copy (Fig. S2A).

**Figure 2.**
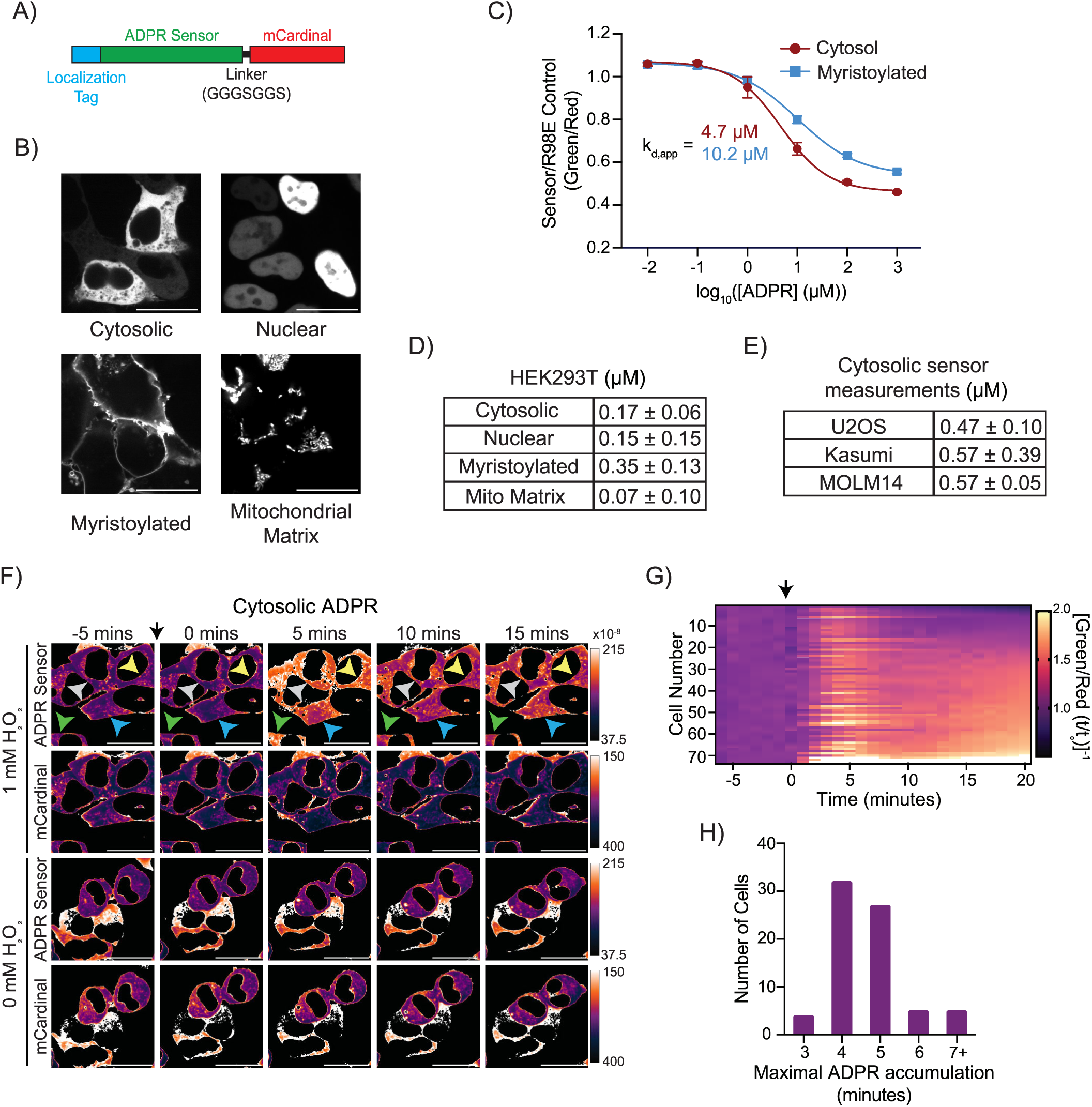
ADP-ribose concentration can be quantified in live cells. **A)** Biosensor cassette for green/red ratiometric measurements in cells. **B)** Localized dual color ADPR sensor-mCardinal expression in stably transduced live HEK293T cells, scale bar 20 µm. **C)** Standard curves measured with flow cytometry using cells expressing localized ADPR sensor-mCardinal and normalized to parallel green/red controls, mean ± SEM, *n* = 3 - 4. **D)** Intrapolated concentration for (2’deoxy/phospho-)ADPR in nuclear, cytosol, myristorylated, and mitochondrial matrix compartments in HEK293T cells, mean ± SEM, *n* = 7 - 20 samples. **E)** Estimated cytosolic concentration of (2’deoxy/phospho-)ADPR in different cell lines based on ratiometric intrapolation with the HEK293T standard curve, mean ± SEM, *n* = 7 – 19 samples. **F)** Representative time-lapsed images of ADPR sensor-mCardinal stably expressed in live HEK293T cells at 34°C; arrow, addition of H_2_O_2_ (1 mM). Green and red channels are depicted separately, and image intensities were inverted and pseudo-colored for viewing, scale bar 20 µm. **G)** Trajectories of ratiometric sensor readouts from individual cells analyzed by live confocal imaging, *n =* 73 cells; arrow, addition of H_2_O_2_ (1 mM). **H)** Distribution of cells based on time to maximal ADPR sensor response.

We generated individual HEK293T lines where the sensor was localized to the cytosol, nucleus, membrane (myristoylated), and mitochondrial matrix (Fig. 2B) and used these lines to calibrate the sensors in cells. We acutely permeabilized with saponin and equilibrated the lines with a range of ADPR concentrations (Fig. 2C, S2B). Analyses were restricted to the subpopulation that both retained sensor fluorescence and showed internalization of propidium iodide (Fig. S2E) indicating equilibration with extracellular small molecules. Apart from the myristolylated sensor, the addition of localization tags did not significantly alter responsiveness of the sensors (Fig. S2C), and so we calibrated both myristoylated and cytosolic sensors. We found the range of sensor measurements in cells was 100 nM to 100 μM, with binding constants ^Cyto^K_d,app_ ∼4.7 μM [95% confidence interval (CI), 3.73 to 5.88] and ^Myr^K_d,app_ ∼10.2 μM [95% CI, 8.24 to 12.90]. With the calibration curve, we intrapolated the concentrations of measured ligand at different subcellular compartments in HEK293T cells (Fig. 2D); all measurements were < 500 nM. We further estimated basal ADPR levels across different cell lines based on the HEK293T curve and all estimates were < 1 µM (Fig. 2E); AML lines were estimated to have slightly higher basal ADPR levels between 500 - 600 nM.

### Single-cell measurements reveal heterogenous nature of ADPR accumulation

A limited number of approaches have been validated to modulate intracellular ADPR, and so we used H_2_O_2_-induced stress as an established approach to characterize the cytosolic sensor in cells (49–52). We observed a mean maximal accumulation of free ADPR at 5 minutes post-treatment and dissipation starting at 10 minutes post-treatment (Fig. 2F).

To determine the heterogeneity of responses to H_2_O_2_-induced stress, we analyzed ADPR accumulation in individual cells and found that single-cell measurements generally mirrored the dynamics of bulk measurements (Fig. 2G, 2H, S2D). We observed and quantified heterogeneity across individual cells with regards to the amplitude and timing of maximal sensor fluorescence response (Fig. 2G, 2H, S2D). The variation was independent of sensor expression (Fig. S2E) and the R98E control did not show significant variability (Fig. S2F, S2G), suggesting that the variability indeed reflected heterogeneity for intracellular ADPR accumulation. Together, the data support the use of sensors for both bulk and single-cell measurements.

### Intracellular ADPR fluctuations are distinct from NAD^+^ levels

We determined that two different doses of exogenous H_2_O_2_ resulted in transient accumulations of cytosolic, nuclear, and membrane ADPR with similar dynamics but distinct maxima (Fig. 3A, S3A-B); no ADPR signal was detected with paired negative control sensors R98E (Fig. S3C-E). We found that H_2_O_2_-induced ADPR depended on PARP1/2 and PARG activities, as the signal was ablated by Olaparib and PDD00017273 treatment, respectively (Fig. 3B-C, S3F-G). Together the data indicated that free ADPR was generated by dismantling of PARP1/2-dependent poly-ADPr chains by PARG.

**Figure 3.**
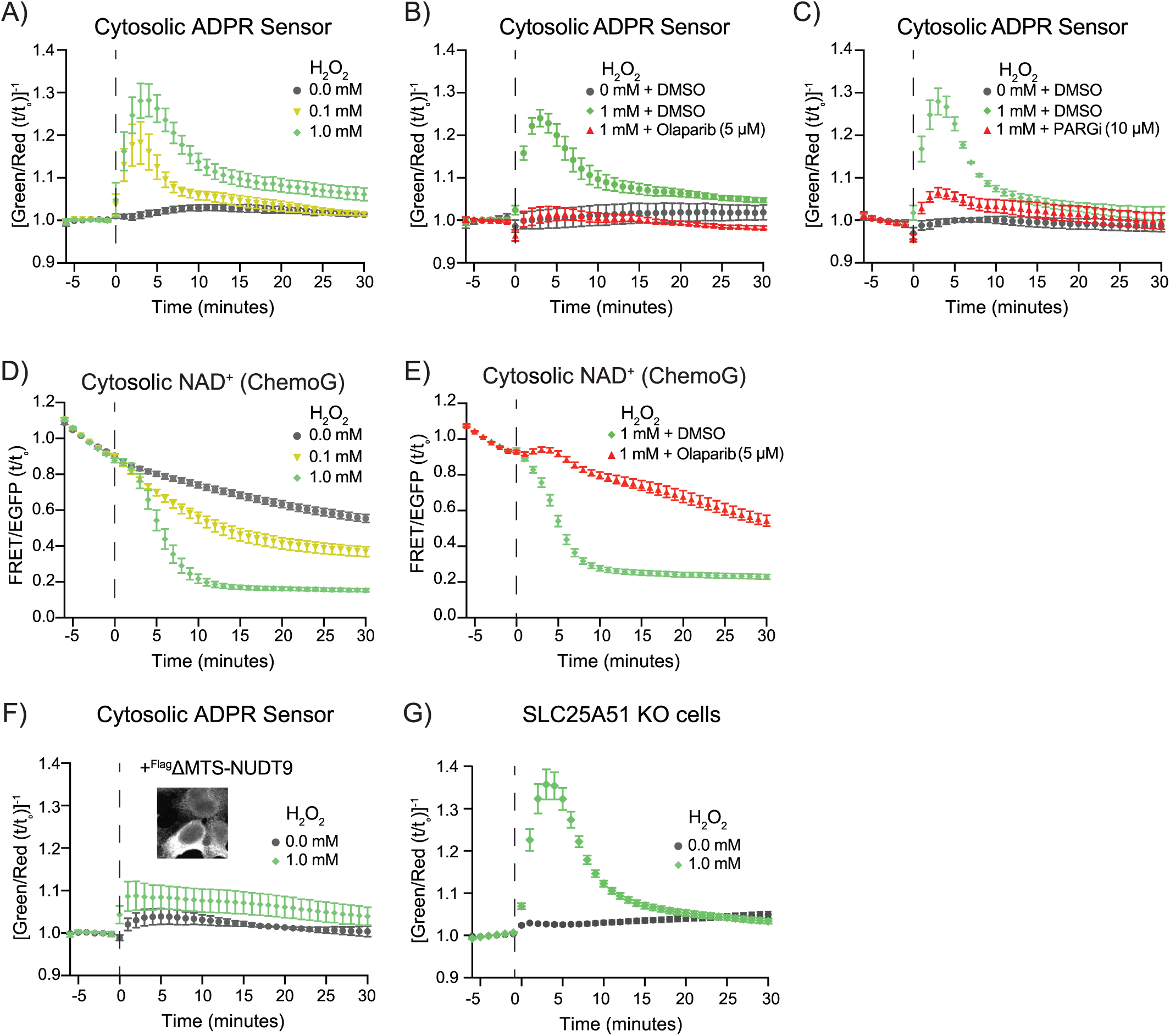
ADPR dynamics are distinct from NAD^+^ fluctuations. **A)** Ratiometric measurements from stably expressed cytosolic ADPR sensor-mCardinal in HEK293T cells at 34°C; dashed line, addition of H_2_O_2_. Data (imaging) are mean ± SEM, *n* = 3 experiments each containing 750 - 1000 cells monitored over time with 10x lens. **B)** Treatment with PARP1/2 inhibitor, olaparib (5 µM) added at *t* = 0 min, *n* = 3 experiments each containing 750 - 1000 cells monitored over time with 10x lens. **C)** Treatment with PARG inhibitor, PDD00017273 (10 µM) added at *t* = 0 min, *n* = 3 experiments each containing 750 - 1000 cells monitored over time with 10x lens. **D)** Cytosolic NAD^+^ levels measured in parallel with stably expressed ChemoG-NAD_JF635_ biosensor, *n* = 9 - 12 experiments each containing 30 - 50 cells monitored over time with 100x lens. **E)** NAD^+^ levels with olaparib (5 µM) added at *t* = 0 min, *n* = 11 - 12 experiments each containing 30 - 50 cells monitored over time with 100x lens. **F)** Effect of ectopic co-expression of cytosolic NUDT9 on cytosolic ADPR sensor, *n* = 3 experiments each containing 750 - 1000 cells monitored over time with 10x lens. **G)** Response of ADPR sensor (cytosolic) in cells with depleted mitochondrial NAD^+^, *n* = 9 experiments each containing 750 - 1000 cells monitored over time with 10x lens.

To compare the dynamics between ADPR accumulation and NAD^+^-consumption, we monitored cytosolic NAD^+^ levels in parallel using a FRET-based ChemoG-NAD biosensor. While the ADPR maxima had peaked at 5 minutes, we interestingly observed that NAD^+^ continued to deplete until 15 – 20 minutes post-treatment and that its concentration remained low (Fig. 3D). The dynamics of NAD^+^ depletion was distinct from any photobleaching observed with the ChemoG-NAD biosensor. The H_2_O_2_-dependent loss of NAD^+^ was prevented by olaparib treatment, supporting that PARP activity was dominantly consuming NAD^+^ (Fig. 3E). Together, the data indicated distinct trajectories following H_2_O_2_-induced stress between free NAD^+^ and free ADPR concentrations.

We next co-expressed a cytosolic-localized NUDT9 enzyme that lacked its N-terminal mitochondrial targeting sequence, ΔMTS-NUDT9 (Fig. 3F, 3G). NUDT9, K_M,(ADPR)_ = 3 μM (53), was previously used to suppress ADPR mediated TRPM2 activation (11). We found that co-expression of ΔMTS-NUDT9 ablated H_2_O_2_-induced ADPR (Fig. 3H) and had no significant effects on the R98E control (Fig. S3H). NUDT9 activity is selective for linear forms of ADPR over cyclic-ADPR, confirming that the sensor was monitoring linear ADPR. This is also the first direct support that Nudix activity can regulate ADPR levels in cells. Notably, the sensor further revealed that intracellular ADPR was robustly induced in cells lacking mitochondrial NAD^+^ transporter SLC25A51 (Fig. 3I, S3I). The HEK293 SLC25A51 KO line (54) was previously shown to have depleted mitochondrial NAD^+^ concentrations, thus the data suggest that H_2_O_2_ production of ADPR did not depend on high levels of mitochondrial NAD^+^.

### A threshold level of intracellular ADPR is required for TRPM2 activation

We next determined whether the observed burst of intracellular ADPR could signal to TRPM2 and mediate influx of extracellular calcium. We used a landing pad HEK293T cell line (55) to integrate at single copy a cassette that co-expressed calcium sensor jGCaMP7s (56) and TRPM2. We found that 1 mM of exogenous H_2_O_2_ was sufficient to trigger calcium influx (Fig. 4A). The timing of channel opening corresponded to peak ADPR levels, and calcium influx was ablated with block of PARP activity (Fig. 4B). Notably, while 100 µM exogenous H_2_O_2_ treatment produced ADPR, the accumulation was less and not sufficient to activate TRPM2. This indicated there exists a threshold concentration of intracellular ADPR required to activate TRPM2 in cells. With 1 mM H_2_O_2_ treatment and channel opening, we observed a second influx of calcium starting at approximately 20 minutes post-treatment that did not directly correspond to the ADPR accumulation. Collectively, the biosensor data revealed a threshold level of ADPR required for TRPM2 activation and demonstrated that proxy measurements such as NAD^+^ consumption and calcium influx were inexact approximations for ADPR fluctuations.

**Figure 4.**
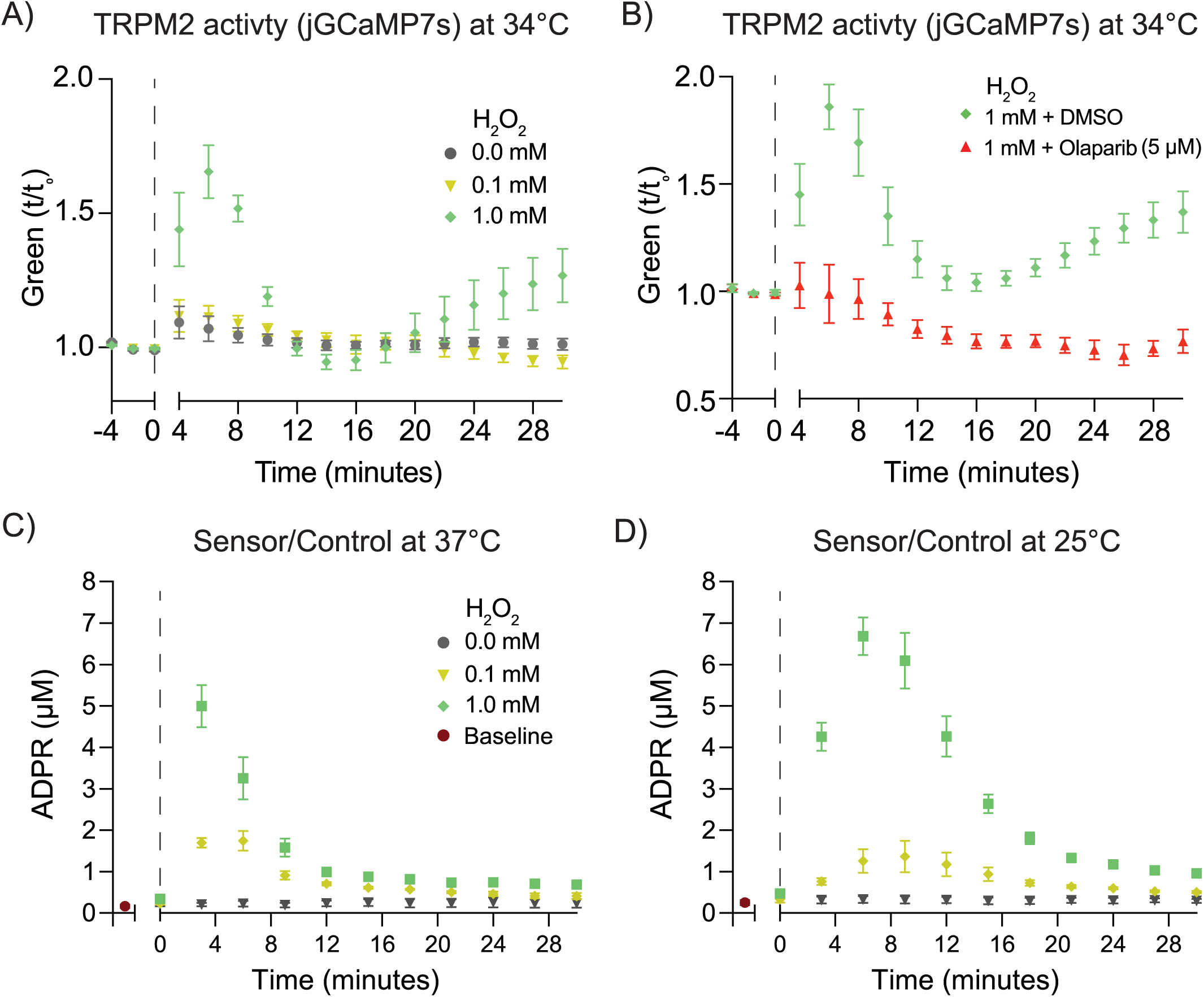
A threshold level of intracellular ADPR is required for TRPM2 activation. **A)** Intracellular calcium measured with jGCaMP7s-IRES- hTRPM2 expressed via single-copy landing pad in HEK293T cells, dashed line, addition of H_2_O_2_. Data (imaging) are mean ± SEM, *n* = 4 - 5 experiments each containing 750 - 1000 cells monitored over time with 10x lens. **B)** Intracellular calcium with treatment with PARP1/2 inhibitor, olaparib (5 µM) added at *t* = 0 min, *n* = 4 - 5 experiments each containing 750 - 1000 cells monitored over time with 10x lens. **C)** Quantified intracellular ADPR at 37°C using control-normalized measurements with the cytosolic sensor, obtained with the same flow cytometer used for the calibration curve in Fig. 2C. Baseline measurement (maroon) prior to addition of H_2_O_2_ (dashed line). Data (cytometer) are mean ± SEM, *n* = 3 experiments. **D)** Quantified intracellular ADPR at 25°C, similar to above.

We next quantified the different concentrations of intracellular ADPR produced. At 37°C, treatment with 1 mM H_2_O_2_ resulted in peak accumulation of 5.00 ± 0.89 μM cytosolic ADPR, which is ∼30-fold higher than pretreatment levels (Fig. 4C). The lower dose (0.1 mM H_2_O_2_) resulted in 1.85 ± 0.25 μM accumulation and was insufficient to activate TRPM2 channels. Therefore we estimate a threshold for activation > 2 μM in cells.

Given the temperature-sensitive nature of TRPM2 activation, we determined how temperature affected H_2_O_2_-induced ADPR by examining resulting levels at 37°C (Fig. 4C) compared to 25°C (Fig. 4D). We observed that the cooler temperature appeared to slow down production and turnover of ADPR. This resulted in ∼2x longer time to peak accumulation, a slightly wider peak, and slightly elevated final ADPR accumulation of 6.68 µM ± 0.8 µM.

## Discussion

We report and characterize a new genetically encoded ratiometric fluorescent biosensor to monitor and quantify ADPR fluctuations in cells (Fig. 1). The biosensor is also sensitive to other physiologically relevant and canonical TRPM2 agonists and Nudix enzyme substrates like 2’-deoxy-ADPR, 2’/3’-O-acetyl-ADPR, and phospho-ADPR (18, 44–47). We localized it to different parts of the cell and obtained the first direct measurements of intracellular ADPR as both bulk and single-cell measurements (Fig. 2).

With the biosensor, we uncovered that intracellular ADPR signaling was transient and tunable, and determined that ADPR was produced and accumulated in response to PARP1/2 and PARG activity. Accumulation of the signaling metabolite was proportional to amount of H_2_O_2_ treatment and depended on PARG-dependent release of ADPR subunits from PAR chains.

Because the maximal accumulation depended on the amount of H_2_O_2_, ADPR concentrations likely reflected the amount of induced damage. The data further suggest that PARG activity was not limiting in the cell and that ADPR biosensor measurements might be useful for reflecting the dynamics of endogenous PARG activity.

ADPR loss occurred with comparable kinetics to its production in our measurements between 25°C - 37°C. We have not determined the mechanism responsible for turning over ADPR, but it is likely a Nudix-family enzyme (57, 58); ectopic expression of NUDT9 was sufficient to block accumulation of ADPR (Fig. 3F). Cells with depleted mitochondrial NAD^+^ levels were not impeded in cytosolic ADPR accumulation (Fig. 3G). This is consistent with H_2_O_2_ treatment activating non-mitochondrial PARP/PARG activity but challenges an earlier interpretation about a mitochondrial source of ADPR (11).

Intriguingly we observed that NAD^+^ consumption continued at a steady rate after ADPR accumulation had reached its peak and slowed down (Fig. 3A, D). A comparison of ∼2 µM vs ∼5 µM ADPR accumulation showed that the curves reached their peaks and turned over at comparable rates (Fig. 3A). So even if ADP-hydroxymethyl-pyrrolidinediol (ADP-HPD) is an amino-analogue of ADPR that can block PARG activity (59), the similar kinetics indicate that product inhibition of PARG was not a major contributor. The effect could instead reflect increased Nudix enzyme activity while PARG activity is being restrained. Other possibilities include that the later NAD^+^ consumption reflected a non-PARP activity or that there may exist a subset of PAR chains simply inaccessible to PARG activity. This will require further study.

By calibrating the biosensor, we were able to obtain the first measurements of intracellular ADPR and found that basal levels were < 1 µM in all tested cell lines. We found two cell lines derived from acute myeloid leukemia that had slightly elevated basal levels of ADPR ∼ 570 nM (Fig. 2E). Stimulation of ADPR production resulted in transient ADPR accumulation representing as much as a 30-fold increase. Notably, we observed that an ADPR accumulation ∼4 µM, but not ∼2 µM, resulted in channel activation of TRPM2 at low expression (Fig. 4). These values are consistent with previously measured ADPR concentrations for physiological channel opening (7, 18, 22, 26, 60). Moreover, we have directly shown that the fluctuation in ADPR levels coincided with the timing of channel opening. Together the data make a strong case demonstrating that ADPR fluctuation is a determining factor for TRPM2 opening under physiological conditions.

We found that temperature did not significantly alter the ultimate amount of accumulated ADPR, but a lowered temperature slowed the rate of ADPR production and turnover (Fig. 4). A limitation of the work is that we did not formally measure ADPR accumulation between 37°C and 42°C due to lack of sensitive temperature control on the calibrated cytometry instrument.

Nevertheless, the data collectively indicate that threshold ADPR concentrations, and not rate of production, is the driving factor for channel opening and this is consistent with TRPM2 being responsible for integrating environmental signals. Additionally, we observed two sequential and separable calcium influx events following H_2_O_2_ treatment. This aligned with some previous reports (15, 24, 25) but differed from other previously observed sustained influxes (2, 11, 61). We suspect the differences may depend on the relative density and expression of TRPM2 in the different experimental systems. In this work, TRPM2 was expressed at single-copy in HEK293T, and the relatively lower density of TRPM2 channels in these cells may have limited sensitization and sustainment of a calcium rise.

Notes on effective use of the ADPR biosensor:

1. Too much expression of the biosensor in cells can buffer ADPR and attenuate its detection. We recommend low or single copy expression of the sensor.
2. ADPR changes can be compartmentalized, and we recommend localizing the sensor at the appropriate subcellular location.
3. pH and temperature can impact measurement values as reported. We recommend the non-binding R98E control for normalization.

## Materials and Methods

### Cells

HEK293T (ATCC: CRL-3216) cells were cultured at 37°C with 5% CO_2_ in Dulbecco’s modified Eagle’s medium (DMEM, Sigma-Aldrich, Cat# D6429) containing glucose (4.5 g/L), sodium pyruvate (1 mM), L-glutamine (4 mM) and supplemented with 10% Fetal Bovine Serum (FBS, Biowest, Cat# S1620), penicillin/streptomycin (1x) and 25 mM HEPES buffer (pH 7.4).

### Lentivirus

500,000 HEK293T cells/well were seeded in 6-well TC-treated plates (Fisherbrand, Cat# FB012927) and transfected the next day using polyethylenimine (PEI) at a ratio of 5:1 PEI:DNA µg. The transfection mixture was prepared in Opti-MEM (Gibco, Cat# 31985-070) and comprised of 20 μg PEI and a plasmid mixture of 1 μg psPAX2, 1 μg pMD2.G, and 2 μg of either pLenti- NES-Flag-HA-ADPR Sensor-mCardinal-IRES-Puro, pLenti- NES-Flag-HA-R98E-mCardinal-IRES-Puro, pLenti- NLS-Flag-HA-ADPR Sensor-mCardinal-IRES-Puro, pLenti- NLS-Flag-HA-R98E-mCardinal-IRES-Puro, pLenti- Myr-ADPR Sensor-mCardinal-IRES-Puro, pLenti- Myr-R98E-mCardinal-IRES-Puro, pLenti-4x(cox8)-Flag-HA-ADPR Sensor-mCardinal-IRES-Puro, pLenti-4x(cox8)-Flag-HA-R98E-mCardinal-IRES-Puro, pLenti-N-Flag-HA-GAW-IRES-Puro, pLenti-N-Flag-HA-Nudt9(59 - 350 amino acids)-IRES-Puro or pLenti-N-Flag-HA-ChemoG-NAD-IRES-Puro plasmids. Supernatant that contained viral particles was collected 72 hours post-transfection and filtered through a 0.45 μm PVDF filter (Millex, Cat# SLHVR33RS). Five hundred microliters of viral supernatant were added to a cell suspension of HEK293T cells (125,000 cells/mL) in a 6-well plate and incubated for 2 days. Transduced cells were either sorted using FACS with a SONY MA900 or selected with puromycin (2 μg/mL) or blasticidin (20 μg/mL) and cultured with selection for 2 and 6 days respectively. Transduced cells were maintained in selection media until all non-transduced control cells succumbed to antibiotic selection in parallel. Stable expression was verified with Western Blotting and live cell imaging.

Generation of HEK293T-iCasp9-LLH-IB cells with attB-GCaMP7s-IRES-TRPM2 integration. 500,000 HEK293T-iCasp9-LLH-IB cells were seeded in 6-well TC-treated plate and transfected next day using PEI:DNA ratio of 5:1 diluted in Opti-MEM. The DNA mix for transfection included 1 μg pCAG NLS-HA Bxb1 and 1 μg attB-GCaMP7s-IRES-TRPM2. 48 hrs post-transfection, cells were selected in media containing 10 nM AP1903 and 1 mM doxycycline for 2 days. The selection was deemed sufficient when the control cells succumbed completely. The selection media was replaced with fresh media containing 1 mM doxycycline and the cells were allowed to recover.

### Plasmids

2x HiFi builder assembly master mix (NEB #E2621S) was used to assemble the following plasmids: pLenti-NES-Flag-HA-ADPR Sensor-mCardinal-IRES-Puro, pLenti-NLS-Flag-HA-ADPR Sensor-mCardinal-IRES-Puro, pLenti-4x(cox8)-Flag-HA-ADPR Sensor-mCardinal-IRES-Puro, pET15b-His-ADPR Sensor, pET15b-His-ADPR Sensor-mCardinal, pLenti-Flag-HA- Nudt9(59 - 350 amino acids)-IRES-Puro and attB-GCaMP7s-IRES-TRPM2. Q5 site-directed mutagenesis kit (NEB #E0554S) was used to generate mutations in the above plasmids and to obtain R98E control variant.

### Protein Purification

*E. coli* BL21 DE3 competent cells were transformed with pET15b-His-ADPR Sensor or pET15b-His-R98E plasmids and plated on LB Agar (50 µg/mL kanamycin, Kan). A single colony was picked in 5 mL LB media (50 µg/mL Kan) and grown for 16 hours shaking at 225 rpm at 37°C. The culture was diluted 1:50 in 250 mL LB Kan media and induced using 1 mM IPTG once an OD_600_ of 0.6 was achieved. Induced cultures were grown at room temperature for 24 hours. Cultures were centrifuged at 4,000 x g for 10 minutes at 4°C and cell pellet was washed in 5 mL equilibration buffer (20 mM Tris HCl, 200 mM NaCl, pH 7.5). Cells were collected by centrifugation (4,000 x g for 10 minutes at 4°C) and the cell pellet was resuspended and lysed in 40 mL lysis buffer (20 mM Tris HCl, 200 mM NaCl, 0.1% Triton-X-100 v/v, 1 mM PMSF, 10 mM Imidazole, pH 7.5) for 30 minutes, rotating at 4°C. Cells were then sonicated using Fisherbrand™ Q705 Sonicator with a 19 mm probe on ice (Amplitude 75, processing time: 1 minute and 30 second, on time: 15 seconds, off time: 45 seconds). Lysate was clarified by centrifugation at 12,000 x g for 30 minutes at 4°C before being filtered through a 0.45 μm PVDF filter (Millex, Cat# SLHVR33RS). A 10 mL disposable gravity column was loaded with 2 mL slurry of Ni-NTA Superflow Resin (Cat# EM70691-4) and washed with 10 mL equilibration buffer followed by 10 mL lysis buffer. The filtered lysate was passed through the gravity flow column twice. The column was washed with 10 mL wash buffer 1 (20 mM Tris-HCl, 200 mM NaCl, 10 mM Imidazole, pH 7.5) and 10 mL wash buffer 2 (20mM Tris-HCl, 200mM NaCl, 25mM Imidazole, pH 7.5). This was followed by the elution in 10 mL elution buffer (20 mM Tris-HCl, 200 mM NaCl, 200 mM Imidazole, pH 7.5) and eluate was collected as 1 mL fractions. The three fractions displaying the brightest fluorescence were combined for further processing. Dialysis was performed using Slide-A-Lyzer™ G3 Dialysis Cassettes (Cat# A52966). Cassettes were floated in 250 mL dialysis buffer (100 mM Tris-HCl pH 7.4, 150 mM NaCl, 0.5 mM DTT, 100 μM PMSF, 1 mM EDTA, 30% glycerol v/v) while stirring for three cycles 2-hour each and an overnight dialysis step with buffer refreshed between each step. Purified protein was resolved using SDS PAGE followed by Coomassie staining. The quantification was done using Bradford assay along with BSA standards.

### Fluorimeter Measurements

ADPR sensor or R98E control recombinant protein (125 nM) was mixed with the ligand at indicated concentrations in fluorescence assay buffer (100 mM Tris-HCl pH 7.4, 150 mM NaCl and 1x Pierce™ Protease Inhibitor Tablets, EDTA-free Cat# A32965)). The fluorescence was measured using Horiba fluorimeter at 520 nm emission following excitation scans from 400-510 nm, as well as with 500 nm excitation and emission scans from 515-600 nm. Tested ligands included ADPR (Sigma-Aldrich, Cat# A0752, 10 μM), 2’-deoxy ADPR (BioLog, Cat# D227, 10 μM), O-Acetyl ADPR (Santa Cruz Biotechnology, Cat# sc-481663, 10 μM), phospho ADPR (Biolog, Cat# A190, 10 μM), 2’-cyclic ADPR (Axxora Marketplace, Cat# BLG-C406-005,10 μM), 3’-cyclic ADPR (Axxora Marketplace, Cat# BLG-C404-005,10 μM), cyclic ADPR (Biolog, Cat# C005, 10 μM), NaAD (Sigma-Aldrich, Cat# N4256, 10 μM), NAD^+^ (Sigma-Aldrich, Cat# N1636, 10 μM), NADH (Roche, Cat# 10128023001, 10 μM), NaADP^+^ (Biolog, Cat# N018, 10 μM), R5P (Sigma-Aldrich, Cat# R7750, 10 μM), AMP (Sigma-Aldrich, Cat# A1752, 500 μM), ADP (Sigma-Aldrich, Cat# 2754, 500 μM), and ATP (Sigma-Aldrich, Cat# 9062, 500 μM).

### Stopped Flow Kinetics Measurements

Stopped flow kinetic experiments were performed using a KinTek AutoSF-120 instrument with a circulating water bath kept at 20°C. The ADPR Sensor (final concentration 125 nM) in fluorescence assay buffer (100 mM Tris-HCl pH 7.4, 150 mM NaCl and 1x Pierce™ Protease Inhibitor Tablets, EDTA-free) was mixed with ADPR at indicated concentrations to start the reaction. A laser light source was used to excite at 515 nm, and emission was monitored with a 565 ± 12 nm bandpass filter (Semrock). The stopped flow traces shown are an average of at least 6 individual traces. KinTek Explorer software version 11 was used to analyze the data as described earlier (62). The parameters *k_1_* and *k_-_*_1_ were obtained by fitting the data to the proposed mechanism: 𝐸 + 𝑆 ⇌ 𝐸𝑆. For equation based analysis of the stopped flow data, traces were fit initially using the analytical (Afit) function of KinTek Explorer to a single exponential function: 𝑦 = 𝐴_O_ + 𝐴_1_(1 − 𝑒^-bt^), where A_0_ is the starting fluorescence, A_1_ is the amplitude of fluorescence change, b is the decay rate, and t is time. Subsequent mechanism-based fitting was achieved using the numerical integration routines in KinTek Explorer.

### In cell Calibration Curve

HEK293T cells were permeabilized with 0.004% saponin (Sigma-Aldrich, Cat# 84510) and 3 μM propidium iodide to first demonstrate degree of permeabilization. To generate a standard curve, HEK293T cells stably expressing cytosolic ADPR sensor and R98E control were trypsinized and 300,000 cells were collected for each measurement. The cells were washed with DPBS and replaced with 100 μL of calcium measurement buffer (140 mM NaCl, 5 mM KCl, 1 mM CaCl2, 1 mM MgSO4, 1 mM NaH2PO4, 20 mM HEPES, and 5.5 mM glucose, pH 7.4). 100 μL of a mixture containing saponin and the indicated ADPR concentration were equilibrated at room temperature for 30 minutes prior to measuring the sensor fluorescence. The final curve was taken using the ratio of the ADPR sensor/R98E control. The values from three biological replicates were used to fit a four-parameter sigmoidal model against ADPR concentration on GraphPad v10.6.0. Using the GraphPad equation, Y=Bottom + (X^Hill^ ^slope^)*(Top-Bottom)/(X^Hill^ ^slope^ + EC50^Hill^ ^Slope^), values from a H_2_O_2_ dosage experiment were interpolated to calculate ADPR concentrations that were produced in response.

### Live-Cell Microscope Imaging Using 10x Lens

HEK293T cells stably expressing the ADPR Sensor or the R98E control were seeded at a density of 250,000 cells/mL in a 24-well TC-treated glass bottom plate (Cellvis, Cat# P24-1.5H-N). HEK293T-iCasp9-LLH-IB cells with attB-GCaMP7s-IRES-TRPM2 integration were seeded similarly after 48 hours induction using 1 mM doxycycline. Prior to imaging, the media was replaced with 200 μL Fluorobrite DMEM (Gibco, Cat# A18967-01). The cells were allowed to equilibrate to new conditions for at least 30 mins prior to imaging. Cell imaging was done in an environment-controlled chamber with 5% CO_2_ at 34°C using OKO-cage system. After imaging for baseline fluorescence, 200 μL of 2x indicated treatment, including H_2_O_2_ (Sigma Aldrich, Cat# 216763), olaparib (LC Laboratories, Cat# O-9201) or PDD00017273 (Cayman Chemicals, Cat# 19511) was added and imaged for indicated time. Cells were imaged using a SOLA SM Light Engine, Photometrics Prime 95B SCMOS digital camera, PLAN ACHROMAT 10x Objective, NA 0.25, WD 10.6 MM. The instrument was automated using a motorized xy-stage for frame positioning and z-axis control to adjust the focus for clear imaging. Multichannel images were captured with an IX3 ET-EGFP/FITC/Cy2 and IX3 ET-Cy5 fluorescence filter, with an excitation of 470 ± 40 nm or 520 ± 60 nm, and an emission of 525 ± 50 nm or 700 ± 75 nm, respectively. Each field of image contained approximately 750 - 1000 fluorescent cells for analysis.

### Microscope Image Quantification For Bulk Analysis

All images captured using 10x lens were analyzed using ImageJ2 v2.16.0/1.54p software. Mean fluorescence intensity was noted from each frame. Background intensities from each frame were subtracted using the thresholding function, and remaining mean fluorescence intensity was measured. Images were minimally processed and involved threshold subtraction for each frame followed with mean fluorescence intensity quantification. The ratiometric sensor measurement was obtained by dividing the ADPR sensor or R98E control fluorescence with mCardinal fluorescence. Data was normalized by dividing each value by the average of baseline measurements for GCaMP7s (Green) and ADPR sensor or R98E control (Green/Red) taken from time 0 to 5 minutes.

### NAD^+^ Monitoring using ChemoG-NAD_JF635_ Sensor

HEK293T cells stably expressing ChemoG-NAD were seeded at 100000/mL in a 24-well glass bottom plate and 24 hours later the media was replaced with fresh media containing 200 nM JF635 dye for overnight incubation. Next day, the media was replaced with 200 μL Fluorobrite DMEM 30 minutes before the imaging. Cell imaging was performed in an OKO-cage system with controlled environment at 34°C and 5% CO_2_. Cells were imaged using UAPO 100XTIRF OBJ, NA 1.49, WD 0.1MM, W/CC lens and a high-powered spinning disk laser system. ChemoG-NAD_JF635_ multichannel images were taken using the excitation wavelength of 488 nm, with fluorescence emission at 525 ± 50 nm and FRET emission at 617 ± 73 nm. We took baseline fluorescence measurements before the addition of the indicated treatments at t = 0 mins. Image analysis was performed using ImageJ where the threshold for the image was set and was subtracted from mean fluorescence intensity for EGFP and FRET. The ChemoG-NAD_JF635_ FRET was calculated by dividing FRET fluorescence with EGFP fluorescence. Each field of view contained approximately 30 – 50 cells.

### Confocal Imaging and Single Cell Analysis of ADPR Sensor using a 100x Lens

HEK293T cells expressing the ADPR Sensor or the R98E control were seeded at a density of 250,000 cells/mL in a 24-well glass bottom plate. Prior to imaging, the media was replaced with 200 μL Fluorobrite DMEM. The cells were allowed to equilibrate to new conditions for at least 30 mins prior to imaging. Cell imaging was done in an environment-controlled chamber with 5% CO_2_ at 34°C using OKO-cage system. For single-cell analysis, imaging was done using a UAPO 100XTIRF OBJ, NA 1.49, WD 0.1MM, W/CC lens and a high-powered spinning disk laser system. We took multichannel images for ADPR sensor and R98E Control using excitation wavelengths of 488 nm and 561 nm, with their respective emission at 525 ± 50 nm and 617 ± 73 nm. All images captured using 100x lens were analyzed using ImageJ2 v2.16.0/1.54p software. Each individual cell was enclosed in a region of interest and followed over the experiment time. Individual experiment’s field of view image contained approximately 5 - 20 fluorescent cells.

The threshold function was used to determine the background for each image and was subtracted from mean fluorescence intensity values. Ratiometric fluorescence values for ADPR Sensor and R98E control were calculated by dividing the sensor fluorescence with mCardinal. For determining the localization of the sensor, the cell nucleus was stained using 20 mM Hoechst 33342 (ThermoScientific, Cat# 62249) diluted 1:2000 in DPBS and incubated at room temperature for 10 minutes, protected from light. The cells were then washed three times with DPBS following which the media was replaced with 200 μL of Fluorobrite DMEM for imaging, with an excitation wavelength of 405 nm and emission wavelength of 525 ± 50 nm.

### Representative Time-Lapsed Images of ADPR Sensor-mCardinal in Live HEK293T Cells

Threshold was applied to create a binary mask for each color in every image. The original image was multiplied to the binary mask in ImageJ using image calculator option, and a reciprocal was taken using the math option in ImageJ. We then selected GEM LUT and image was further smoothened. The whole image was cropped to show representative cells.

### Flow Cytometry

Sensor fluorescence was measured by analyzing 10,000 fluorescent cells using flow cytometry with NovoCyte Flow Cytometer 3000. The fluorescence was measured for ADPR Sensor and R98E at excitation 488 nm, emission 530 ± 30 nm and the fluorescence for mCardinal was measured at excitation 561 nm, emission 660 ± 20. Standard gating was applied to exclude debris and cell doublets. Flow cytometry data was analyzed using FlowJo v10.

### Western Blot

500,000 HEK293T cells were lysed in 100 μL 2x Laemmli sample buffer (Tris pH 6.8, glycerol, DTT, 10% SDS, bromophenol blue), followed by an incubation at 95°C for 5 minutes. 15 μL of cell lysate was resolved by SDS-PAGE using NUPAGE 4-12% Bis-Tris protein gel (Invitrogen) with MOPS running buffer. Following transfer to a 0.45 μm nitrocellulose membrane, the membrane was blocked with 5% BSA in pH 7.4 Tris-buffered saline with 0.1% (v/v) Tween 20 (TBST). The primary antibodies were incubated overnight at 4°C in TBST containing 1% BSA, followed by 3 washes of 5 minutes each with TBST. Secondary antibodies in TBST containing 1% BSA were incubated for an hour at room temperature followed by 3 washes of 5 min each with TBST. Images were taken on an Odyssey CLx Imager. The antibody dilutions used: anti-FLAG (Sigma-Aldrich, Cat# F1804, 1:3000), anti-Tubulin (Sigma-Aldrich, Cat# T9026, 1:3000); and anti-mouse IgG H&L Alexa Fluor 680 (Invitrogen, Cat# A10038, 1:10000).

### Immunostaining Imaging

22x22 mm glass coverslips coated with 25 μg/mL Poly-L-Lysine (Sigma-Aldrich, Cat# A-005-M) were seeded with 300,000 mammalian cells in a TC-treated 6-well plate. For cell fixation, the media was removed and cells were fixed in 4% paraformaldehyde (Electron Microscopy Sciences, Cat# 15710) and 4% sucrose in DPBS for 10 minutes. Fixed cells were washed with DPBS and transferred to a humidifying chamber to prevent evaporation. Cells were blocked and permeabilized in 2% BSA, 5% normal goat serum, 0.3% Triton-X in DPBS for 10 minutes.

Primary antibody was incubated overnight at 4°C in the blocking buffer. After the removal of the primary antibody, secondary antibody was placed and incubated at room temperature for one hour. Coverslips were mounted face-down on 15 μL of Vectashield Vibrance Anti-Fade Mounting Medium with DAPI (Fisher Scientific, Cat# NC1601054) on microscope slides. After curing at room temperature for one hour, the sides were sealed with nail polish. Multichannel images were taken using an excitation wavelength of 405 nm and 561 nm, with their respective emission wavelength of 460 ± 50 nm and 617 ± 73 nm. Antibodies and dilutions used were as follows: rabbit anti-FLAG (Cell Signaling Technology, Cat# 14793, 1:500) and anti-rabbit IgG Highly Cross-Absorbed Alexa Fluor 561 (Invitrogen Cat# A-11036, 1:1000).

### Statistical Analysis

The data were analyzed and plotted using GraphPad Prism 10 v10.5.0. Statistical significance was determined using unpaired two-sided Student’s t-test for two samples, ordinary one-way ANOVA with post-hoc Dunnett’s or Tukey’s test for more than two samples, and Mann-Whitney U test using a false discovery rate (FDR) of 0.01% and two-stage step-up method for non-normal data. *P*-values under 0.05 were deemed significant with \**p* < 0.05, \*\**p* < 0.01 and \*\*\**p* < 0.001.

For global fit analyses using KinTek Explorer software (63), residuals were computed as the difference between each datapoint and the mean. Normal distribution around zero indicated an accurate fit of model to the data. Standard deviation was calculated from the data using the equation 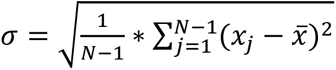, where 𝑥*_j_*− 𝑥̅ are residuals.

Confidence contour plots in Fig. 1F for k_on_ and k_off_ values, as well as for fluorescence scaling factors *a* and *b*, were calculated using the FitSpace (64) function of the KinTek Explorer software. The following output observable expression was used in data fitting: a*(E+(b*ES)), where *a* is the scaling factor for the initial measured fluorescence and *b* is the scaling factor for the fractional change in signal upon ligand binding. Confidence contour plots showed that calculated parameters were well-constrained by the data (Fig. 1F).

## Data Availability

All data supporting the findings in this study are provided in the figures and Supplementary files. Full view images and source data have been deposited to the Texas Data Repository, Cambronne XA Lab Dataverse, https://doi.org/10.18738/T8/DBSGWU.

## Acknowledgements

We thank Cambronne lab members, as well as Jonathan Smailys, Jessie Zhang, Ralf Fliegert, Tessa Schwarzer, and Andreas Guse for critical discussions. We thank Andrés Jara-Oseguera for the jGCaMP7s calcium sensor in the landing pad plasmid and Can Cenik for the landing pad HEK293T cell line. We thank Mason Galliver and Jessica Taylor for technical support. This work was supported by awards from the NIH R01CA272490, R35GM152218, and the Pew Charitable Trust.

## Author Contributions

S.G. and X.A.C. conceptualized this work; S.G., V.N., S.N.L. T.L.D., K.A.J. and X.A.C. designed research; S.G., V.N., S.N.L., T.L.D., W.Y, C.F., H.C., Y.M., L.C., N.R. and K.D. performed research; S.G., V.N., S.N.L., T.L.D., W.Y., K.A.J, and X.A.C analyzed data; S.G., V.N. and X.A.C. wrote the paper; all authors reviewed the paper; X.A.C. supervised the project and acquired the funds for this work.

## Competing Interest Statement

S.G and X.A.C are inventors on a US patent application filed by The University of Texas at Austin covering the technology described here.

---

**Figure S1.**
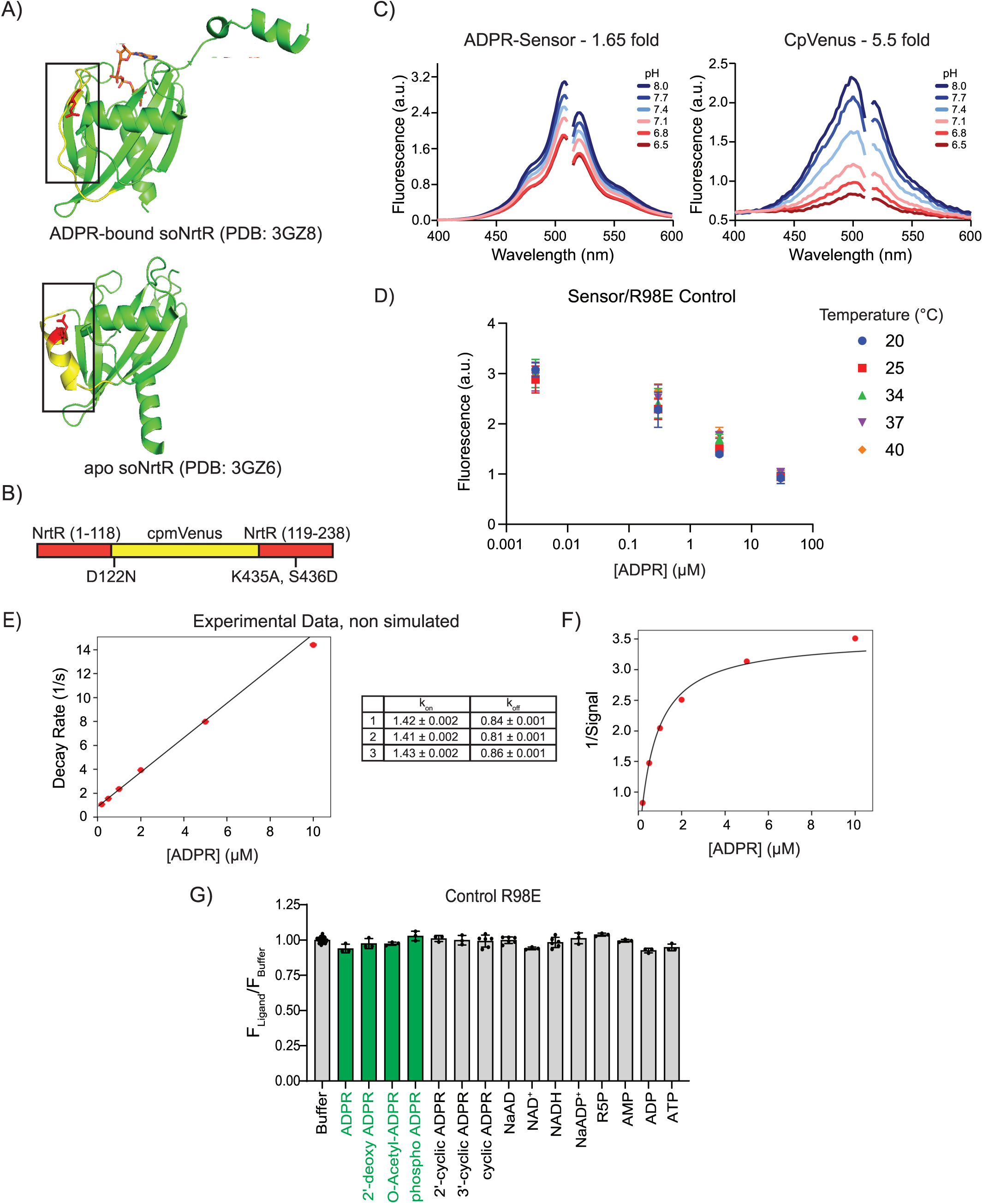
Supplemental figure panels accompanying main text Figure 1. *Design and in-vitro characterization of ADPR Sensor and R98E Control*. **A)** Schematic of ADPR biosensor design. **B)** Conformation of α-helix 2 when unbound (bottom, PDB 3GZ6) compared to ADPR-bound (top, PDB 3GZ8) (44). **C)** pH effects on fluorescence brightness of the ADPR biosensor, left, compared to another fluorescent protein circularly permutated (cp)Venus, right. **D)** Dose-dependent response of ADPR biosensor when normalized to the R98E control at indicated temperatures, *n* = 3. **E)** Representative experimental dataset used for global fit simulation of sensor kinetics. Calculated experimental values across 3 replicates are indicated. **F)** Representative experimental dataset represented as 1/fluorescence, compared to ADPR concentration. **G)** Effect of ADPR and indicated molecules of R98E sensor control. All ligands were tested at 10 µM except AMP/ADP/ATP at 500 µM, reflecting their expected physiological intracellular concentrations, *n =* 54 without and *n =* 3 - 6 with ligands.

**Figure S2.**
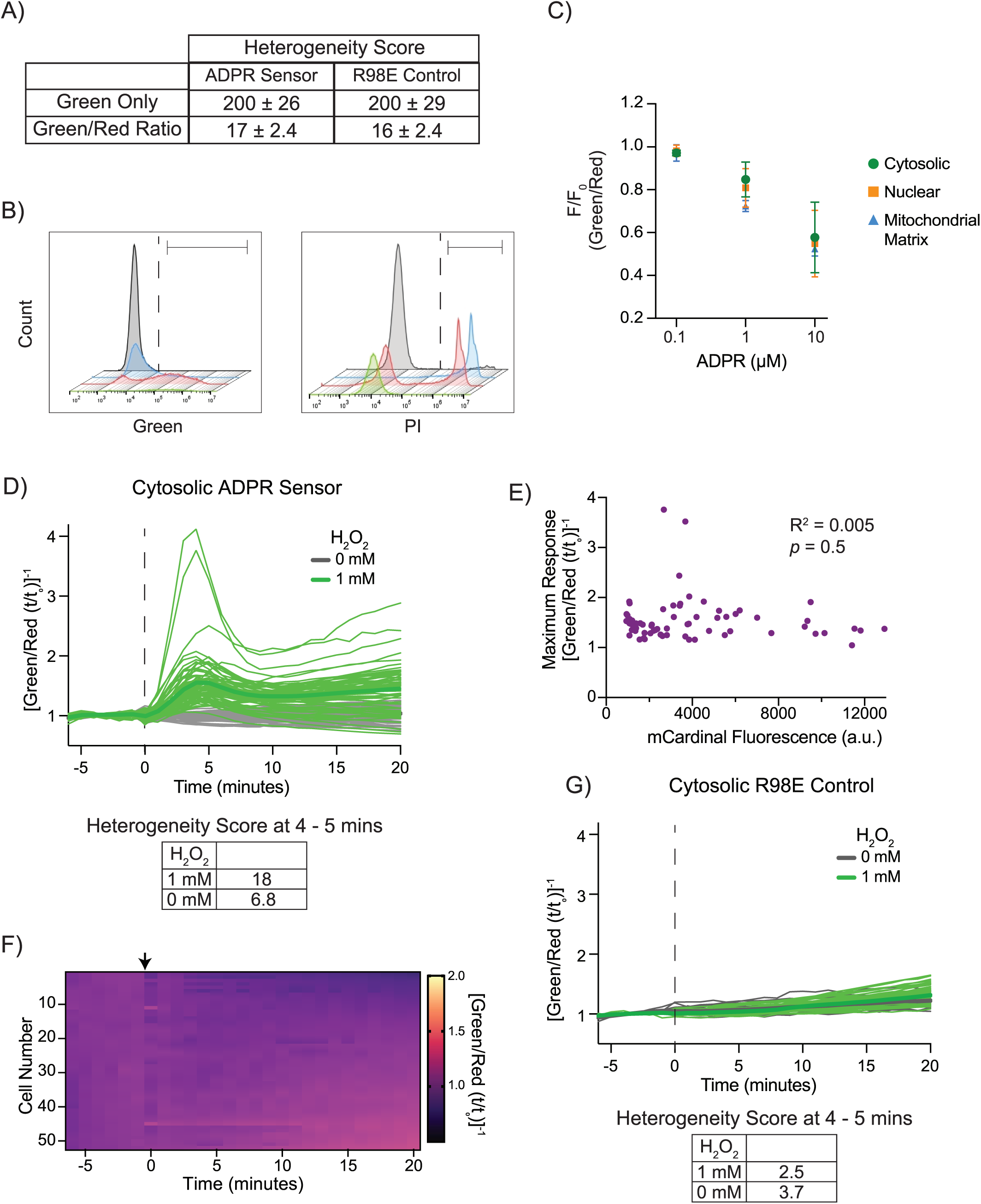
Supplemental figure panels accompanying main text Figure 2. *Sensor normalization, identification of population to use for calibration, single cell measurements*. **A)** Quantitation using a heterogeneity score of the distribution of basal biosensor readouts across stably expressed cell lines when measured as green-only intensity compared to ratiometric green/red fluorescence. Heterogeneity Score = (interquartile range of the distribution)/ (geometric mean of distribution). Data mean scores ± SD from, *n* = 40 experiments each containing 10000 cells. **B)** Three subpopulations can be resolved after cell permeabilization and equilibration with propidium iodide (PI) based on retention of biosensor green fluorescence and internalization of red PI, relative to untreated cells (grey). Only the “red” population was defined by green fluorescence and internalization of red PI, and only readouts from this subpopulation were included in the calibration curve, n = 10000 cells. **C)** Biosensor variants with cytosolic, nuclear, and mitochondrial localization tags responded similarly to ADPR. Data (fluorimeter) mean ± SD, *n* = 3. **D)** Individual traces of the cytosolic ADPR-mCardinal sensor across cells shown in 2G (n= 52 - 73), following addition of buffer or H_2_O_2_ (dashed line). Mean traces are bolded green and grey, and variability in the maxima between 4 - 5 minutes is indicated by heterogeneity scores below. **E)** Maximum amplitude from ADPR-mCardinal sensor readouts in individual cells compared to mCardinal fluorescence in same cell (n= 73), *p* = 0.5 **F)** Trajectories of ratiometric R98E control measurements from individual cells analyzed by live confocal imaging, *n =* 27 - 68; arrow, addition of H_2_O_2_ (1 mM). **G)** Individual traces of the cytosolic R98E control across cells (n= 27 - 68) following addition of buffer or H_2_O_2_ (dashed line). Mean traces are bolded green and grey; calculated heterogeneity scores from maxima between 4 - 5 minutes.

**Figure S3.**
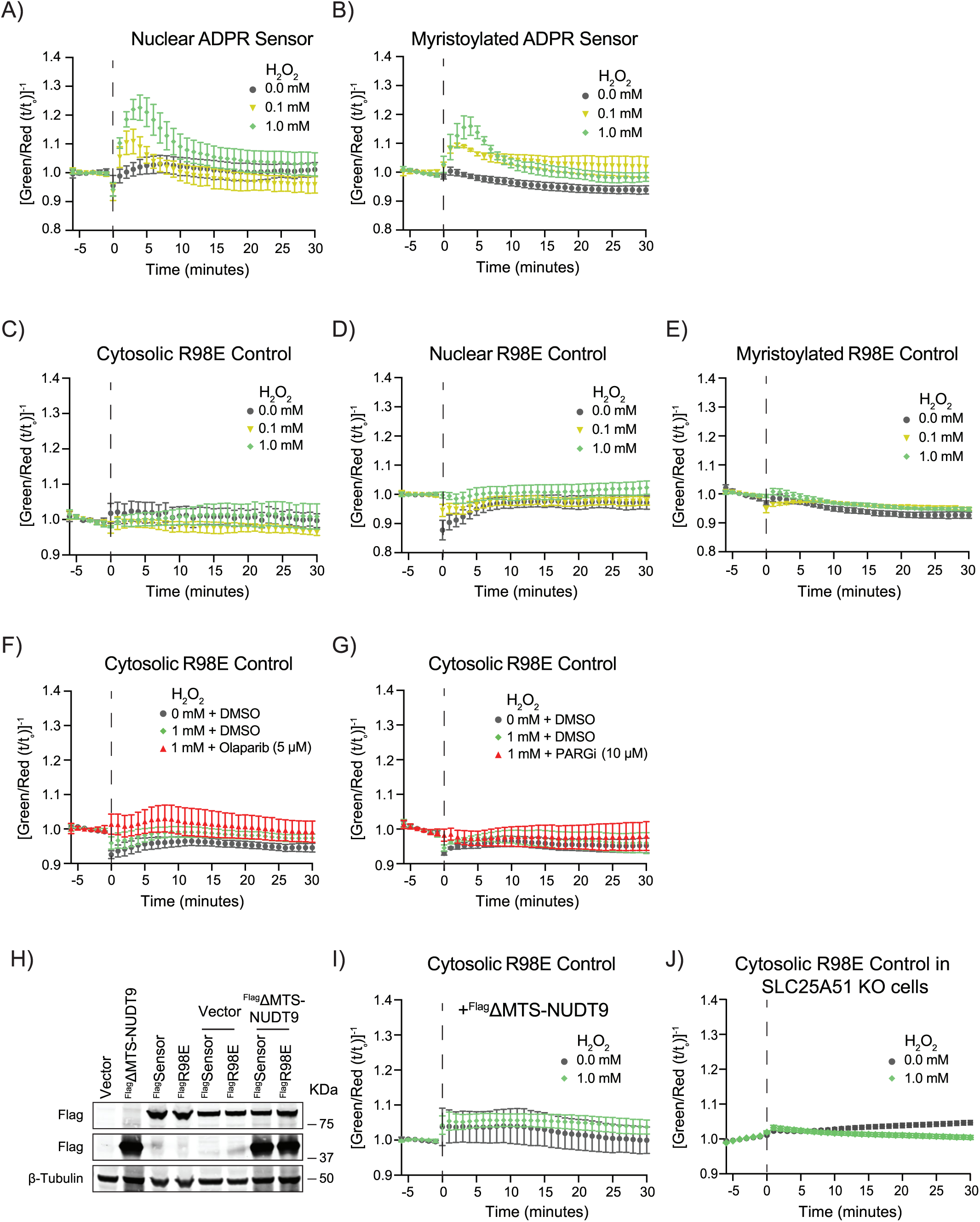
Supplemental figure panels accompanying main text Figure 3. *Reponses of localized sensor, responses of R98E controls, expression of ^Flag^ΔMTS-NUDT9*. **A)** Ratiometric measurements from stably expressed nuclear ADPR sensor-mCardinal in HEK293T cells; dashed line, addition of H_2_O_2_. Data (imaging, 34°C) are mean ± SEM, *n* = 3 experiments each containing 750 - 1000 cells monitored over time with 10x lens. **B)** Response from myristoylated ADPR sensor-mCardinal in HEK293T cells; Data (imaging, 34°C) are mean ± SEM, *n* = 3 experiments each containing 750 - 1000 cells monitored over time with 10x lens. **C-G)** Response from localized R98E-mCardinal controls as indicated, and with either Olaparib or PDD00017273 added at *t* = 0 min as indicated. Data (imaging, 34°C) are mean ± SEM, *n* = 3 experiments each containing 750 - 1000 cells monitored over time with 10x lens. **H)** Western blot depicting ectopic expression of ^Flag^ADPR Sensor (MW 88.3 kDa) , ^Flag^R98E Control (MW 88.3 kDa), or ^Flag^ΔMTS-NUDT9 (MW 37.5 kDa) individually or co-expressed as indicated. Loading control, β-tubulin (MW 50.5 kDa). **I)** Response from the R98E-mCardinal control to co-expression of ^Flag^ΔMTS-NUDT9 *n* = 3 experiments each containing 750 - 1000 cells monitored over time with 10x lens. **J)** Response from the R98E-mCardinal control in SLC25A51 KO HEK293 cells. *n* = 3 experiments each containing 750 - 1000 cells monitored over time with 10x lens.

## Notes

### Competing Interest Statement

The University of Texas at Austin has filed a patent application covering the technology described here.

### Summary of Updates

Updates to this version: - Figure 4 has been updated - Minor additional text edits

## References

1. B. Vilar, C.-H. Tan, P. A. McNaughton, Heat detection by the TRPM2 ion channel. Nature 584, E5–E12 (2020).

2. Z. Zhong, et al., TRPM2 links oxidative stress to NLRP3 inflammasome activation. Nat. Commun. 4, 1611 (2013).

3. C.-H. Tan, P. A. McNaughton, The TRPM2 ion channel is required for sensitivity to warmth. Nature 536, 460–463 (2016).

4. K. Uchida, et al., Lack of TRPM2 Impaired Insulin Secretion and Glucose Metabolisms in Mice. Diabetes 60, 119–126 (2011).

5. I. Lange, et al., TRPM2 Functions as a Lysosomal Ca ^2+^ -Release Channel in β Cells. Sci. Signal. 2, 23 (2009).

6. A. Di, et al., The redox-sensitive cation channel TRPM2 modulates phagocyte ROS production and inflammation. Nat. Immunol. 13, 29–34 (2012).

7. I. Lange, R. Penner, A. Fleig, A. Beck, Synergistic regulation of endogenous TRPM2 channels by adenine dinucleotides in primary human neutrophils. Cell Calcium 44, 604–615 (2008).

8. H.-W. Liu, et al., Bilirubin gates the TRPM2 channel as a direct agonist to exacerbate ischemic brain damage. Neuron 111, 1609–1625.e6 (2023).

9. K. Song, et al., The TRPM2 channel is a hypothalamic heat sensor that limits fever and can drive hypothermia. Science (1979). 353, 1393–1398 (2016).

10. K. Togashi, et al., TRPM2 activation by cyclic ADP-ribose at body temperature is involved in insulin secretion. EMBO J. 25, 1804–1815 (2006).

11. A.-L. Perraud, et al., Accumulation of Free ADP-ribose from Mitochondria Mediates Oxidative Stress-induced Gating of TRPM2 Cation Channels. Journal of Biological Chemistry 280, 6138–6148 (2005).

12. S. Chen, et al., Transient receptor potential ion channel TRPM2 promotes AML proliferation and survival through modulation of mitochondrial function, ROS, and autophagy. Cell Death Dis. 11, 247 (2020).

13. M. Gershkovitz, et al., TRPM2 Mediates Neutrophil Killing of Disseminated Tumor Cells. Cancer Res. 78, 2680–2690 (2018).

14. K. Essuman, et al., The SARM1 Toll/Interleukin-1 Receptor Domain Possesses Intrinsic NAD + Cleavage Activity that Promotes Pathological Axonal Degeneration. Neuron 93, 1334–1343.e5 (2017).

15. S. Yamamoto, et al., TRPM2-mediated Ca2+ influx induces chemokine production in monocytes that aggravates inflammatory neutrophil infiltration. Nat. Med. 14, 738–747 (2008).

16. A. H. Guse, Enzymology of Ca2+-Mobilizing Second Messengers Derived from NAD: From NAD Glycohydrolases to (Dual) NADPH Oxidases. Cells 12, 675 (2023).

17. B. Tóth, L. Csanády, Identification of Direct and Indirect Effectors of the Transient Receptor Potential Melastatin 2 (TRPM2) Cation Channel. Journal of Biological Chemistry 285, 30091–30102 (2010).

18. B. Tóth, I. Iordanov, L. Csanády, Ruling out pyridine dinucleotides as true TRPM2 channel activators reveals novel direct agonist ADP-ribose-2′-phosphate. Journal of General Physiology 145, 419–430 (2015).

19. W. M. Riekehr, et al., cADPR Does Not Activate TRPM2. Int. J. Mol. Sci. 23, 3163 (2022).

20. R. Fliegert, W. M. Riekehr, A. H. Guse, Does Cyclic ADP-Ribose (cADPR) Activate the Non-selective Cation Channel TRPM2? Front. Immunol. 11, 548219 (2020).

21. K. Nagamine, et al., Molecular Cloning of a Novel Putative Ca2+Channel Protein (TRPC7) Highly Expressed in Brain. Genomics 54, 124–131 (1998).

22. A.-L. Perraud, et al., ADP-ribose gating of the calcium-permeable LTRPC2 channel revealed by Nudix motif homology. Nature 411, 595–599 (2001).

23. Y. Sano, et al., Immunocyte Ca2+ Influx System Mediated by LTRPC2. Science (1979). 293, 1327–1330 (2001).

24. Y. Hara, et al., LTRPC2 Ca2+-Permeable Channel Activated by Changes in Redox Status Confers Susceptibility to Cell Death. Mol. Cell 9, 163–173 (2002).

25. E. Wehage, et al., Activation of the Cation Channel Long Transient Receptor Potential Channel 2 (LTRPC2) by Hydrogen Peroxide. Journal of Biological Chemistry 277, 23150–23156 (2002).

26. Á. Bartók, L. Csanády, Dual amplification strategy turns TRPM2 channels into supersensitive central heat detectors. Proceedings of the National Academy of Sciences 119, e2212378119 (2022).

27. B. Bastide, K. Snoeckx, Y. Mounier, ADP-ribose stimulates the calcium release channel RyR1 in skeletal muscle of rat. Biochem. Biophys. Res. Commun. 296, 1267–1271 (2002).

28. A. J. Gustafsson, et al., ADP ribose is an endogenous ligand for the purinergic P2Y1 receptor. Mol. Cell. Endocrinol. 333, 8–19 (2011).

29. C. Huang, et al., Extracellular Adenosine Diphosphate Ribose Mobilizes Intracellular Ca2+ via Purinergic-Dependent Ca2+ Pathways in Rat Pulmonary Artery Smooth Muscle Cells. Cellular Physiology and Biochemistry 37, 2043–2059 (2015).

30. R. H. G. Wright, et al., ADP-ribose–derived nuclear ATP synthesis by NUDIX5 is required for chromatin remodeling. Science (1979). 352, 1221–1225 (2016).

31. H. Qi, R. H. Grace Wright, M. Beato, B. D. Price, The ADP-ribose hydrolase NUDT5 is important for DNA repair. Cell Rep. 41, 111866 (2022).

32. T. V Zharova, A. D. Vinogradov, A competitive inhibition of the mitochondrial NADH-ubiquinone oxidoreductase (Complex I) by ADP-ribose. Biochimica et Biophysica Acta (BBA) - Bioenergetics 1320, 256–264 (1997).

33. V. G. Grivennikova, A. B. Kotlyar, J. S. Karliner, G. Cecchini, A. D. Vinogradov, Redox-Dependent Change of Nucleotide Affinity to the Active Site of the Mammalian Complex I. Biochemistry 46, 10971–10978 (2007).

34. J. A. Birrell, G. Yakovlev, J. Hirst, Reactions of the Flavin Mononucleotide in Complex I: A Combined Mechanism Describes NADH Oxidation Coupled to the Reduction of APAD+, Ferricyanide, or Molecular Oxygen. Biochemistry 48, 12005–12013 (2009).

35. D. Wohlwend, et al., Structures of 3-acetylpyridine adenine dinucleotide and ADP-ribose bound to the electron input module of respiratory complex I. Structure 32, 715–724.e3 (2024).

36. L. Tong, S. Lee, J. M. Denu, Hydrolase Regulates NAD+ Metabolites and Modulates Cellular Redox. Journal of Biological Chemistry 284, 11256–11266 (2009).

37. E. Gattkowski, et al., Analysis of ligand binding and resulting conformational changes in pyrophosphatase NUDT9. 288, 6769–6782 (2021).

38. K. Nowak, et al., Engineering Af1521 improves ADP-ribose binding and identification of ADP-ribosylated proteins. Nat. Commun. 11, 5199 (2020).

39. N. Dani, et al., Combining affinity purification by ADP-ribose-binding *macro* domains with mass spectrometry to define the mammalian ADP-ribosyl proteome. Proceedings of the National Academy of Sciences 106, 4243–4248 (2009).

40. S. Challa, et al., Development and characterization of new tools for detecting poly(ADP-ribose) in vitro and in vivo. Elife 11 (2022).

41. A. Thomas, et al., A Genetically Encoded Sensor for Real-Time Monitoring of Poly-ADP-Ribosylation Dynamics In Vitro and in Cells. ACS Sens. 9, 5246–5252 (2024).

42. E. O. Serebrovskaya, et al., Genetically Encoded Fluorescent Sensor for Poly-ADP-Ribose. Int. J. Mol. Sci. 21, 5004 (2020).

43. D. A. Rodionov, et al., Transcriptional regulation of NAD metabolism in bacteria: NrtR family of Nudix-related regulators. Nucleic Acids Res. 36, 2047–2059 (2008).

44. N. Huang, et al., Structure and Function of an ADP-Ribose-Dependent Transcriptional Regulator of NAD Metabolism. Structure 17, 939–951 (2009).

45. R. Fliegert, et al., 2′-Deoxyadenosine 5′-diphosphoribose is an endogenous TRPM2 superagonist. Nat. Chem. Biol. 13, 1036–1044 (2017).

46. O. Grubisha, et al., Metabolite of SIR2 Reaction Modulates TRPM2 Ion Channel. Journal of Biological Chemistry 281, 14057–14065 (2006).

47. L. A. Rafty, M. T. Schmidt, A.-L. Perraud, A. M. Scharenberg, J. M. Denu, Analysis of O-Acetyl-ADP-ribose as a Target for Nudix ADP-ribose Hydrolases. Journal of Biological Chemistry 277, 47114–47122 (2002).

48. J. Chu, et al., Non-invasive intravital imaging of cellular differentiation with a bright red-excitable fluorescent protein. Nat. Methods 11, 572–578 (2014).

49. A.-L. Perraud, et al., NUDT9, a Member of the Nudix Hydrolase Family, Is an Evolutionarily Conserved Mitochondrial ADP-ribose Pyrophosphatase. Journal of Biological Chemistry 278, 1794–1801 (2003).

50. B. Buelow, Y. Song, A. M. Scharenberg, The Poly(ADP-ribose) Polymerase PARP-1 Is Required for Oxidative Stress-induced TRPM2 Activation in Lymphocytes. Journal of Biological Chemistry 283, 24571–24583 (2008).

51. E. Fonfria, et al., TRPM2 channel opening in response to oxidative stress is dependent on activation of poly(ADP-ribose) polymerase. Br. J. Pharmacol. 143, 186–192 (2004).

52. C. Blenn, P. Wyrsch, J. Bader, M. Bollhalder, F. R. Althaus, Poly(ADP-ribose)glycohydrolase is an upstream regulator of Ca2+ fluxes in oxidative cell death. Cellular and Molecular Life Sciences 68, 1455–1466 (2011).

53. E. Gattkowski, et al., Analysis of ligand binding and resulting conformational changes in pyrophosphatase NUDT9. FEBS J. 288, 6769–6782 (2021).

54. S. Goyal, et al., Dynamics of SLC25A51 reveal preference for oxidized NAD+ and substrate led transport. EMBO Rep. 24, e56596 (2023).

55. K. A. Matreyek, J. J. Stephany, D. M. Fowler, A platform for functional assessment of large variant libraries in mammalian cells. Nucleic Acids Res. 45, e102–e102 (2017).

56. H. Dana, et al., High-performance calcium sensors for imaging activity in neuronal populations and microcompartments. Nat. Methods 16, 649–657 (2019).

57. J. Carreras-Puigvert, et al., A comprehensive structural, biochemical and biological profiling of the human NUDIX hydrolase family. Nat. Commun. 8, 1541 (2017).

58. V. A. Kulikova, A. A. Nikiforov, Role of NUDIX Hydrolases in NAD and ADP-Ribose Metabolism in Mammals. Biochemistry (Moscow) 85, 883–894 (2020).

59. J. T. Slama, et al., Specific Inhibition of Poly(ADP-ribose) Glycohydrolase by Adenosine Diphosphate (Hydroxymethyl)pyrrolidinediol. J. Med. Chem. 38, 389–393 (1995).

60. M. Kolisek, A. Beck, A. Fleig, R. Penner, Cyclic ADP-Ribose and Hydrogen Peroxide Synergize with ADP-Ribose in the Activation of TRPM2 Channels. Mol. Cell 18, 61–69 (2005).

61. R. Fliegert, et al., Ligand-induced activation of human TRPM2 requires the terminal ribose of ADPR and involves Arg1433 and Tyr1349. Biochemical Journal 474, 2159–2175 (2017).

62. K. A. Johnson, Z. B. Simpson, T. Blom, Global Kinetic Explorer: A new computer program for dynamic simulation and fitting of kinetic data. Anal. Biochem. 387, 20–29 (2009).

63. K. A. . Johnson, Kinetic analysis for the new enzymology : using computer simulation to learn kinetics and solve mechanisms. 480 (2019).

64. K. A. Johnson, Z. B. Simpson, T. Blom, FitSpace Explorer: An algorithm to evaluate multidimensional parameter space in fitting kinetic data. Anal. Biochem. 387, 30–41 (2009).

